# Online Transcranial Random Noise stimulation improves perception at high levels of visual white noise

**DOI:** 10.1101/2020.06.22.165969

**Authors:** Michael D. Melnick, Woon Ju Park, Sholei Croom, Shuyi Chen, Lorella Batelli, Ania Busza, Krystel R. Huxlin, Duje Tadin

**Affiliations:** Department of Brain & Cognitive Sciences, University of Rochester, Rochester, NY, USA; Center for Visual Science, University of Rochester, Rochester, NY, USA; FineLab, Human Neuroscience, University of Washington, Seattle, WA, USA; Computational Cognitive Science Group, Brain and Cognitive Sciences, Massachusetts Institute of Technology, Cambridge, MA, USA; Berenson-Allen Center for Noninvasive Brain Stimulation and Department of Neurology, Beth Isrel Deaconess Medical Center, Harvard Medical School, Boston, MA, USA; Center for Neuroscience and Cognitive Systems@UniTn, Istituto Italiano Di Tecnologia, Rovereto, Italy; Department of Psychology, Harvard University, Cambridge, MA, USA; Department of Neurology, University of Rochester, Rochester, NY, USA; Flaum Eye Institute, University of Rochester, Rochester, NY, USA; Department of Neuroscience, University of Rochester, Rochester, NY, USA

**Keywords:** tRNS_1_, perception_2_, stimulation_3_, noise_4_, psychophysics_5_, vision_6_

## Abstract

Transcranial random noise stimulation (tRNS), a relatively recent addition to the field of non-invasive, electrical brain stimulation, has been shown to improve perceptual and cognitive functions across a wide variety of tasks. However, the underlying mechanisms of visual improvements caused by tRNS remain unclear. To study this question, we employed a well-established, equivalent-noise approach, which measures perceptual performance at various levels of external noise and is formalized by the Perceptual Template Model (PTM). This approach has been used extensively to infer the underlying mechanisms behind changes in visual processing, including those from perceptual training, adaptation and attention. Here, we used tRNS during an orientation discrimination task in the presence of increasing quantities of external visual white noise and fit the PTM to gain insights into the effects of tRNS on visual processing. Our results show that tRNS improves visual processing when stimulation is applied during task performance, but only at high levels of external visual white noise—a signature of improved external noise filtering. There were no significant effects of tRNS on task performance after the stimulation period. Of interest, the reported effects of tRNS on visual processing mimic those previously reported for endogenous spatial attention, offering a potential area of investigation for future work.

## Introduction

Transcranial electrical stimulation (tES) is a form of noninvasive neurostimulation that inputs low-current electrical signals into the brain via scalp electrodes. Among the various forms of tES, transcranial direct current stimulation (tDCS) is the most well-established, with interest in its applications increasing steadily (Nitsche & Paulus’ 2000 methods paper has nearly 4,000 citations; Google Scholar, May, 2020). tDCS delivers current in a directional manner with different outcomes depending upon electrode configuration (i.e., cathodal *versus* anodal stimulation). A related method, transcranial alternating current stimulation (tACS), is agnostic to electrode configuration, but requires picking a specific alternating frequency, and is often paired with electroencephalography (EEG) and specific hypotheses regarding oscillatory neural frequencies [e.g., alpha, beta, gamma, theta bands (Kuo & Nitsche, 2012)].

Most recently, Terney and colleagues (2008) described a type of tACS termed transcranial random noise stimulation (tRNS). tRNS employs alternating current with frequencies randomly selected from a defined range (e.g., low-frequency tRNS: 0.1-100Hz, and high-frequency tRNS: 101-640 Hz). The result is white noise in the electrical frequency spectrum within the defined range of frequencies. These parameters give rise to a technique that exposes neurons in the brain to weak, random frequency, oscillating electrical current between the locations of applied electrodes. tRNS has grown in popularity thanks to a number of factors: (1) generally positive results for improving neural processes like perceptual learning (Fertonani et al., 2011, Herpich et al., 2019, Pirulli et al., 2013, van der Groen and Wenderoth, 2016), visual perception (van der Groen and Wenderoth, 2016, Fertonani et al., 2015, Herpich et al., 2019) and cortical excitability (Terney et al., 2008); (2) little to no physical sensation during stimulation, resulting in less discomfort and simpler blinding during experiments, and (3) simpler experimental design than tDCS and tACS, which require choosing cathodal *versus* anodal stimulation or a specific stimulation frequency, respectively. Given these advantages, interest in tRNS has grown; the clearer the underlying mechanisms become, the more useful this technique is likely to become.

Collectively, tES methods have both encouraged and frustrated scientific inquiry, with examples of strong, weak and even null effects. Moreover, we have limited direct evidence about underlying mechanisms driving these results in humans. Considering what is currently known about different tES methodologies, tDCS, tACS and tRNS are all thought to alter cortical excitability in humans (assessed via single-pulse, transcranial magnetic stimulation motor or phosphene thresholds (Herpich et al., 2018, Terney et al., 2008). Importantly, none of these stimulation methods introduce a current large enough to create illusory stimuli such as those resulting from transcranial magnetic stimulation (TMS), such as visual phosphenes or stimulation-induced involuntary movements. In other words, in comparison to the direct signal potentiation induced by TMS, those of tES are considered modulatory. Beyond this general insight however, accounts of how tES affects brain function quickly become complicated. For example, the effects of tES often differ across different brain regions (Nitsche and Paulus, 2000, Kanai et al., 2010, Terney et al., 2008), with anodal tDCS increasing excitability in motor cortex (Nitsche and Paulus, 2000, Nitsche and Paulus, 2001), but generating mixed results when applied over visual cortex (reviewed in Miniussi & Ruzzoli 2013)

Current research suggests different mechanisms of action for different types of tES, with insights increasingly provided by computational models (Bikson et al., 2012, Datta et al., 2012, Edwards et al., 2013, Miranda et al., 2006). These models generally aim to estimate how much current is actually reaching cortex in a typical neurostimulation session in adults, clinical populations or in children (Datta et al., 2012, Truong et al., 2013, Kessler et al., 2013, Minhas et al., 2012, Datta et al., 2010). Homeostatic mechanisms and changes in cellular membrane conductance appear to describe the slow-onset effects of both anodal and cathodal tDCS (Antal and Herrmann, 2016, Bikson et al., 2004, Bindman et al., 1964, Creutzfeldt et al., 1962, Jefferys, 1981). In contrast, induced (neural entrainment) or altered (in frequency or amplitude) neural oscillations appear to account for many effects observed with tACS (Antal and Herrmann, 2016). With respect to tRNS, stochastic resonance has been suggested as a plausible mechanism of action (Fertonani and Miniussi, 2017, van der Groen and Wenderoth, 2016, van der Groen et al., 2018, van der Groen et al., 2019). In their 2016 study, van der Groen & Wenderoth concluded that signals generated by tRNS have similar effects as added visual white noise. This is notable as visual noise can, under certain stimulus condition improve perception via stochastic resonance. Briefly, stochastic resonance is a phenomenon whereby adding noise to subthreshold stimuli can boost performance by raising the overall neural response to the stimulus plus noise combination above a certain threshold. Van der Groen & Wenderoth (2016) showed that tRNS with injected currents ranging from 0 to 1.5mA during a visual, 2-interval, forced-choice task produced inverted U-shaped curves for sub-threshold stimuli that were remarkably similar to stochastic-resonance-induced curves generated when visual white noise was directly added to the stimulus. This exciting result, however, only provided explanation for behavioral improvements resulting from online administration of short bursts of tRNS. It is unclear if stochastic resonance or short bursts of tRNS could explain or elicit the full spectrum of behavioral improvements observed with tRNS (Fertonani and Miniussi, 2017), including longer-term perceptual improvements, such as those seen when tRNS enhances visual perceptual learning (Herpich et al., 2019).

In the present study, we used an equivalent noise paradigm to better understand the mechanisms driving tRNS-induced visual improvements when stimulation is within typical empirical and clinical parameters for duration and current. We probed perceptual changes using a well-established and carefully modeled psychophysical task, where perceptual thresholds were measured as a function of systematically varying external noise levels. This approach assumes that: (1) there is a constant amount of internal noise in the brain, and (2) perceptual performance at an external noise level is influenced by whichever noise (i.e., internal or external) dominates (**Figure 1**). By way of a simplifying analogy, consider a noisy party. In the absence of additional noise, the baseline party noise—corresponding to internal noise—will determine the limits of what one can hear (thick green line, **Figure 1**). Added external noise (e.g., caused by air conditioning; thin orange line, **Figure 1**) will have negligible effects as long as its level is lower than the baseline party noise (**Figure 1**, left box). If the added noise is strong (e.g., a malfunctioning air conditioner), it will start affecting the ability of partygoers to hear each other (**Figure 1**, right box). The negative effects of added noise, however, can be mitigated by an appropriate perceptual template that excludes noise in the incoming signal—a process described as “external noise filtering” in the framework used here. The result is a characteristic threshold-vs-noise function (red line, **Figure 1**) that is flat at low levels of external noise and starts to rise when external noise exceeds internal noise. This framework has been formalized by the Perceptual Template Model or PTM (Dosher & Lu 1998ab; Dosher & Lu 1999a; Lu & Dosher, 1998; Dosher & Lu, 2005; Lu & Dosher, 2008; Zhao et al, 2015) and draws on a long line of research using equivalent noise methodologies (for reviews see Pelli, 1981, Dosher and Lu, 2005). The PTM has been successfully used to explain both how attention enhances visual processing, and improvements caused by perceptual learning (Zhao et al., 2015, Lu and Dosher, 2008, Lu and Dosher, 1998). As such, the PTM is an ideal tool for understanding visual processing changes caused by tRNS.

**Figure 1:**
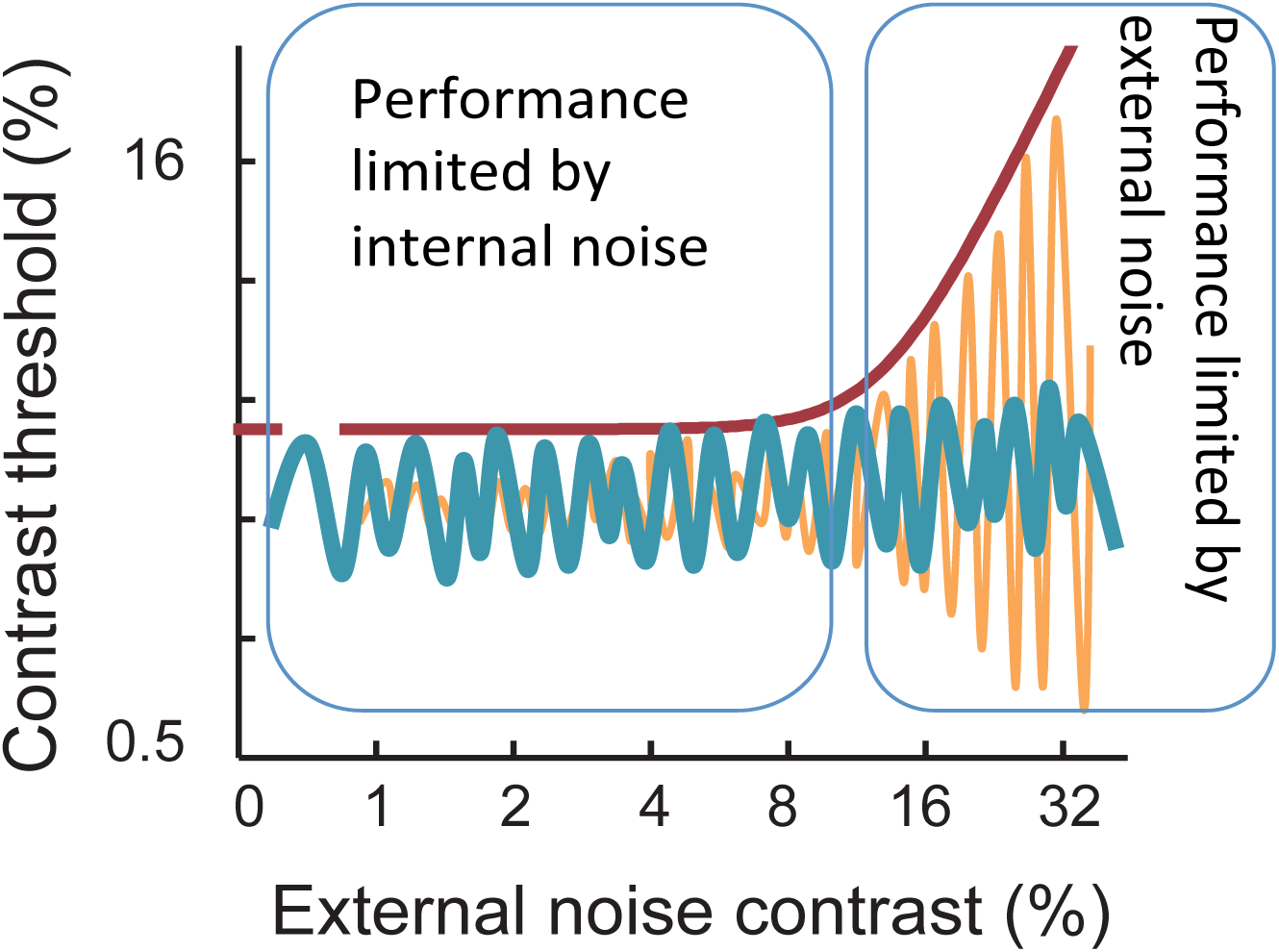
Visual demonstration of key principles underlying equivalent noise paradigms. Example threshold vs. noise curve, where X axis is the percent noise contrast of visual white noise, and the Y axis is the percent contrast threshold of signal stimuli. Performance limitations (in terms of luminance contrast threshold) at low levels of external noise are dominated by internal noise, while performance deficits at high levels of external noise increase linearly with additional external noise. Thick green line: internal system noise; orange line: added external noise; dark red line: perceptual performance.

## Materials and Methods

Ten adults (7 females) were recruited from the undergraduate and graduate populations at the University of Rochester, aged 18-32 years old, with normal or corrected-to-normal vision. Subjects were screened for exclusionary criteria as pertains to electrical stimulation (neural shunts, plates, stents or implants), and informed about the nature and risks of transcranial stimulation. All subjects gave written, informed consent and were paid per hour of participation. Experimental procedures were conducted according to an experimental protocol approved by the University of Rochester’s Research Subjects Review Board and adhered to the tenets of the Declaration of Helsinki.

One subject was ultimately excluded from analysis after fitting normal distributions between subjects at all external noise levels. The fitting revealed that this subject consistently performed at a threshold greater than 2 standard deviations worse than mean performance at all external noise levels, and was affected by the task ceiling performance (100% contrast). As a result, a total of 9 subjects are included in all analyses below, with 4 subjects receiving sham stimulation first, and 5 subjects receiving tRNS stimulation first (see below for more information).

### Experimental Design

Subjects were asked to complete 10 sessions of psychophysical testing, split across 3 consecutive days (**Figure 2**). Day 1 had two intervention-free experimental sessions and was used to induce and assess procedural task-learning. Baseline 1 on Day 1 allowed subjects to gain an understanding of the task and practice performing it at all difficulty levels over a total of 560 trials. Data from Baseline 2, Day 1, was then used as the “true” baseline against which performance collected during subsequent sessions were contrasted. Days 2 and 3 were counterbalanced by subject to assess within-subject effects of tRNS *versus* sham stimulation, and the time course of tRNS-induced behavioral effects.

**Figure 2:**
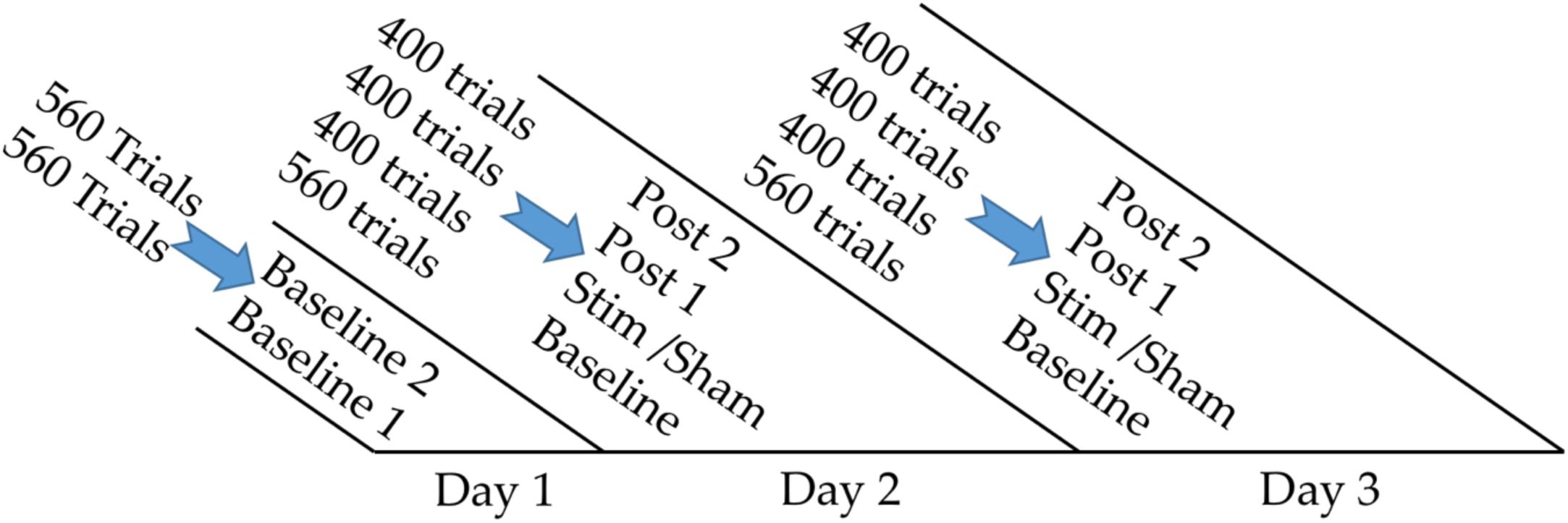
Experimental design. Number of trials performed by all subjects during Days 1-3. Days 2 and 3 were counterbalanced by subject such that half of tested subjects received stimulation on Day 2, and half received stimulation on Day 3. Post 1 was begun 10 minutes after the end of Stim/Sham, or 30 minutes after the beginning of Stim/Sham. Post 2 commenced 10 minutes after the end of Post 1, or a full 60 minutes after the beginning of Stim/Sham.

Subjects were randomized into two groups, with half of them receiving tRNS on Day 2 and sham stimulation on Day 3, and *vice versa* for the other half of the subjects. Brain stimulation (tRNS or sham – see below for details) was administered while subjects performed 400 trials of an orientation discrimination task (see below). This was followed by two 400-trial post-stimulation sessions (Post 1 and Post 2) conducted starting 10 minutes after the end of tRNS/sham sessions (Post 1) and 10 minutes after the end of the Post 1 session (Post 2; **Figure 2**). The resulting session distribution was such that if stimulation began at time 0, Post 1 and Post 2 began ∼30 minutes and ∼60 minutes after the beginning of the stimulation, respectively. In sum, this design allowed us to probe for offline aftereffects of tRNS stimulation which are often observed in other tES methodologies (Pirulli et al, 2013) while the counterbalanced tRNS/sham sessions allowed us to probe for online effects.

### Stimuli

All stimuli were created in MATLAB and displayed using The Psychophysics Toolbox (Brainard, 1997) on a luminance-calibrated CRT monitor (24-inch Sony GDM-FW900 CRT; 1024×640 resolution; 120Hz). Subjects sat comfortably, 77cm from the screen, in a chin-and-forehead rest. Subjects viewed 1.5° diameter, static, non-flickering Gabor stimuli presented centrally and tilted ±12° from vertical, with spatial frequency set to 1 cycle per degree (cpd) (**Figure 3**). Stimulus appearance was preceded by a central fixation cue but fixation was not actively monitored. Subjects responded using the left/right arrow keys to indicate - in this 2-alternative, forced-choice task - whether the Gabor was tilted to the left or right of vertical (**Figure 3**). Gabor stimuli were temporally sandwiched between white noise (**Figure 3**) sampled from a Gaussian distribution that varied in root mean squared contrast in 8 steps, evenly log-distributed between 0 and 33%. This contrast range ensured that we successfully capture the entire threshold *versus* noise (TvN) curve, as confirmed by our pilot work and prior studies in the lab (Park et al., 2017). Rapid sandwiching of the Gabor stimulus between white noise frames allowed for the full, potential, dynamic range of contrast to be used (0 to 100%) while still maintaining the percept of a single Gabor, with varying quantities of noise in the image (Lu and Dosher, 1998, Dosher and Lu, 1999, Lu and Dosher, 2004). With 16.7ms frame duration, this resulted in a total stimulus presentation time of 50ms. Participants received auditory feedback on all trials to indicate the correctness of their responses. Trials did not advance until a response was registered, and subjects were given brief rest periods (30 seconds) every 200 trials to help alleviate fatigue.

**Figure 3:**
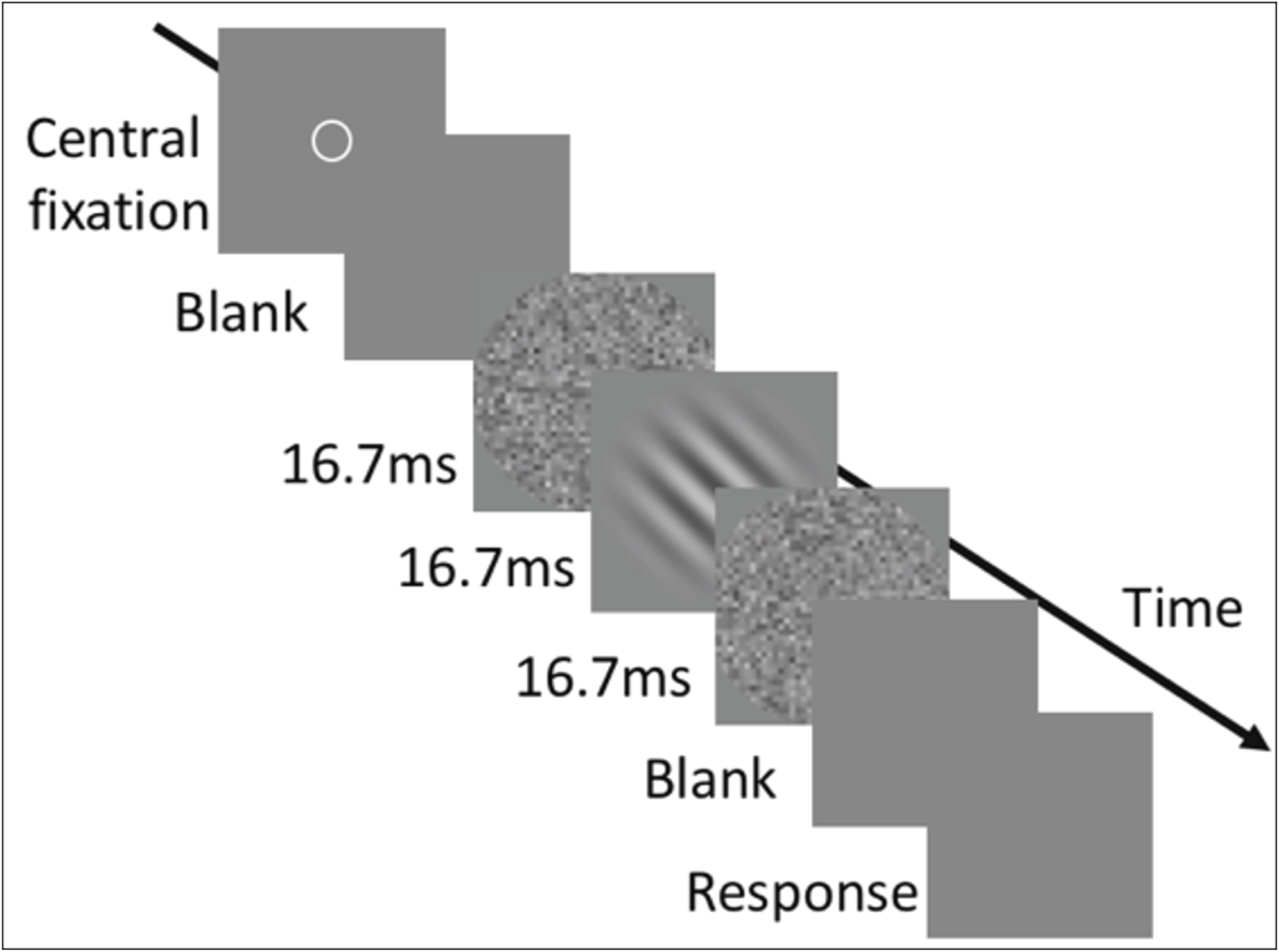
Example trial epoch and stimulation presentation. Central fixation was presented at the onset of each trial. Gabor stimuli were temporally sandwiched between two independent white noise frames of adaptively staircased contrast. Trials did not advance until a subject had responded using keyboard arrows-keys: left for tilted-left, or right for tilted-right. Auditory feedback was provided on all trials.

### tRNS and sham stimulation protocols

All stimulation was delivered using a NeuroConn DC-stimulator PLUS (neuroCare Group, Germany) via 5×7cm sponges soaked in 0.9% NaCl solution. Guided by prior work, tRNS was performed using the high frequency profile (100-640hz) (Herpich et al., 2019, Terney et al., 2008, van der Groen et al., 2019, van der Groen et al., 2018, van der Groen and Wenderoth, 2016). tRNS was applied bilaterally at 2.0mA over occipital cortex at O1 and O2 EEG electrode sites (**Figure 4**). Each subject’s head was measured from inion to occiput and measured for circumference at this height as established in the 10 - 20 EEG system. Centers of electrode sites were then established as 10% of the distance from inion to occiput above and 20% the circumference laterally.

**Figure 4:**
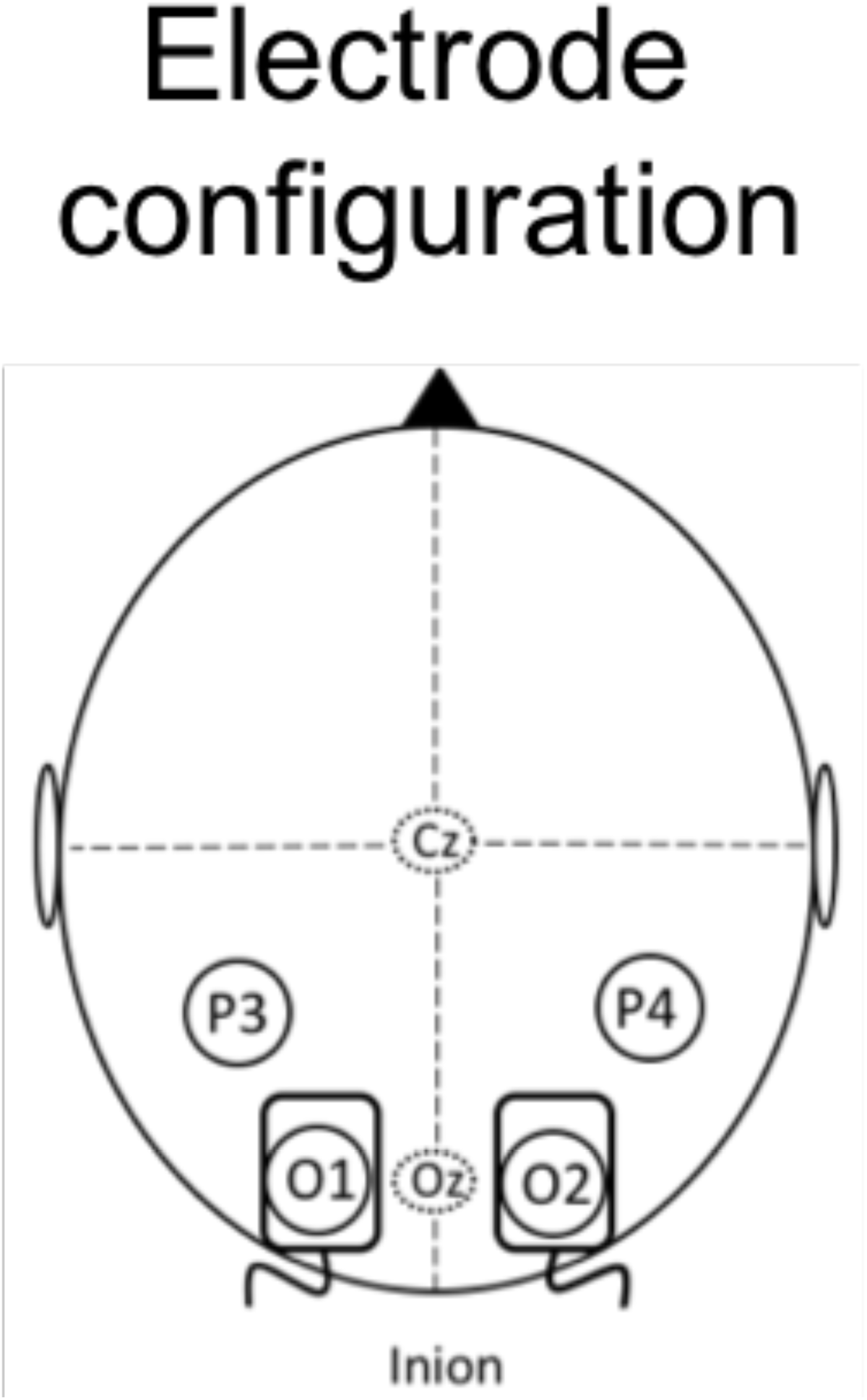
Electrode Montage Configuration for tRNS delivery. The electrode montage configuration used on all subjects based on the 10-20 EEG electrode system. Measurements were acquired for each subject to determine O1 and O2 locations as 10% of the distance from inion to occiput above and 20% the circumference laterally

During stimulation days, stimulation was started just before starting the psychophysical trials. The experimenter verified that subjects felt no discomfort during either sham or tRNS conditions, and as reported previously, tRNS was pain-free and sensationless (Fertonani et al., 2015). On Sham sessions, subjects were set up identically, with an identical time of stimulation. However, current was set to 0mA. This resulted in a behaviorally identical setup for tRNS and sham sessions, including the auditory cues for when the stimulator started and stopped. After stimulation ended, subjects finished their psychophysical session (if they had not already) and were unhooked to take a 10 minutes break before returning for their Post 1 sessions.

### FAST TvN protocol

TvN curves are typically estimated through thousands of trials using the method of constant stimuli (Dosher and Lu, 1998, Dosher and Lu, 2005). However, this is problematic for neurostimulation, which typically lasts only ∼20 minutes (Fertonani et al., 2011, Pirulli et al., 2013, Herpich et al., 2019). To address this issue, we previously developed a parameterized version of the TvN curve using a FAST adaptive staircase method (Park et al., 2017, Vul et al., 2010). This approach employs a single procedure at each difficulty level (71 and 79% correct performance thresholds, corresponding to d’ values of 1.089 and 1.634) that attempts to adaptively estimate the underlying TvN curve across all levels of noise. This results in significantly faster estimation of the TvN curve, as threshold level performance is established as a function of the underlying, estimable TvN parameters (Park et al., 2017, Barbot et al., 2020).

### PTM Analysis

Testing using FAST (Barbot et al., 2020, Park et al., 2017, Vul et al., 2010) did allow us to collect sufficient data within a 20 min brain stimulation session. To analyze our data similarly to earlier work (Dosher and Lu, 1998, Dosher and Lu, 1999, Dosher and Lu, 2005), we pooled data from two FAST procedures yielding 50 trials at each noise level (Park et al, 2017). We then fit individual Weibull psychometric functions (Equation 1) to each noise level using Maximum Likelihood Estimation (MLE):

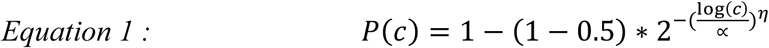

where P corresponds to Percent Correct, c is the stimulus contrast condition, *α* is the 75% psychophysical threshold and η is the slope of the psychometric function. This resulted in 8 psychometric functions per subject per condition, from which 71 and 79% percent correct thresholds were read out.

Conventional PTM fitting was performed on the mean thresholds across subjects by condition. MLE was used to fit the PTM as follows:

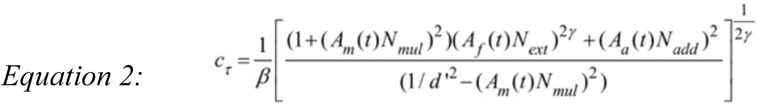

Here, group coefficients are used to distinguish relative differences between each parameter of interest. ***A***_***m***_ represents multiplicative noise, ***A***_***f***_ external noise exclusion, and ***A***_***a***_ additive internal noise. ***γ*** is the fit nonlinearity in the contrast system responsible for multiplicative shifts, and **β** represents the gain on the perceptual template, while ***c***_***τ***_ is the resulting contrast threshold.

Reduced models were fit when comparing conditions by fixing coefficient values. That is, identical quantities of internal, external, and multiplicative noise were used in an attempt to explain two conditions (such as tRNS *versus* sham). This reduced model acted as the null hypothesis for the F-test (Equation 3) comparisons to fuller models that allowed each combination of the 3 coefficients to vary freely (8 combinations in total).

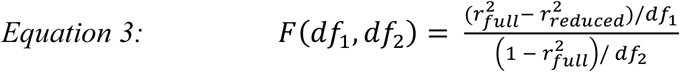

Where *df*_*1*_ = *k*_*full*_ *– k*_*reduced*_ and *df*_*2*_ = *N*_*data*_ – *k*_*full*._ *N*_*data*_ is the number of predicted points (1 for each level of external noise as used here), and *k*_*full*_ and *k*_*reduced*_ are the number of free parameters used in full and reduced models (4 for the reduced models and up to 7 for full). F-tests of full *versus* reduced models were performed with regularization for added parameters, such that added parameters truly increased the explained variance while penalizing for greater free parameters in the model. Mean thresholds across subjects were compared by condition, and only condition comparisons of interest were performed to avoid multiple comparisons.

All in all, we performed three main comparisons: (1) change in baseline performance over subsequent days of testing, (2) online effects of tRNS, comparing online tRNS *versus* online sham stimulation, and (3) offline effects of tRNS, comparing offline tRNS *versus* offline sham stimulation. In each of these comparisons, we first fit the mean contrast threshold across subjects for each external noise level before reading out values for 71 and 79% correct. These average threshold *versus* noise measurements were then compared for the effect of changes in three coefficients in the PTM representing changes in internal additive noise, internal multiplicative noise, and external noise exclusion as detailed above. As is common for PTM studies, we did not find any significant changes in the multiplicative noise parameter (*A*_*m*_), a parameter that is evident in larger changes at higher levels of performance (79 vs. 71% correct). Thus, for data shown below, we present one curve per condition, corresponding to 79% correct performance.

## Results

### Daily change in baseline performance

Before examining the impact of tRNS on visual performance, we first verified that our adaptive method was sensitive enough to pick up the changes in PTM parameters from session to session, which are expected in the context of the well-established phenomenon of visual perceptual learning (Dosher and Lu, 1998, Dosher and Lu, 1999, Lu and Dosher, 2004). To do so, we compared Day 1, Baseline 2 *versus* Day 2, Baseline as the two most comparable sessions, both occurring before any stimulation was delivered. After Day 2, these effects could potentially be confounded by longer-lasting effects of tES, which in some cases has been shown to interfere with perceptual learning (Peters et al., 2013). By using only the second baseline session on Day 1 as a comparator, we minimized the effects of practice-like procedural and task learning that likely dominated the first baseline session.

In comparing Day 1 Baseline 2 *versus* Day 2 Baseline, we saw a significant change in threshold *versus* noise curves with improved performance on Day 2 **(Figure 5)**. The best fitting model indicated an 11.6% improvement in external noise filtering (*F*_*(1,28)*_*=6*.*69, p=0*.*015*) when compared to a reduced model with fixed coefficients for additive internal noise, multiplicative internal noise, and external noise exclusion **(Figure 5, Table 1)**. Perceptual learning studies typically also find an additional effect on reducing additive internal noise (Dosher and Lu, 1998, Dosher and Lu, 2005). Here, the model that had improvements in both internal noise and external noise filtering was marginally significant (*F*_*(2,27)*_*=3*.*23, p=0*.*055*). Note that we restricted our quantitative analysis to the results of PTM fits. Our requirement to collect all data within ∼20 minutes of testing did not give us enough power to analyze individual thresholds for different levels of external noise, since each was based on only 50 trials of data. However, for visualization purposes, we computed changes in performance for each noise level (**Figure 5**). The results, while quite noisy as expected given the number of trials per contrast level, show improvements at all external noise levels. This is a well-established feature of visual perceptual leaning (Dosher and Lu, 1998, Dosher and Lu, 2005) and shows that our approach has sufficient sensitivity to detect expected effects of perceptual learning on perception of noisy stimuli.

**Table 1.**

| Model Noise Variables | $r^2$ | Total Free Parameters | p-value | F |
| --- | --- | --- | --- | --- |
| Base Model: | 0.96 | 3 |  |  |
| Internal Only | 0.96 | 4 | 0.79508 | F(1,28) = 0.068757 |
| External Only | 0.968 | 4 | **0.01518** | F(1,28) = 6.6916 |
| Multiplicative Only | 0.965 | 4 | 0.0563 | F(1,28) = 3.965 |
| Int and Ext | 0.968 | 5 | 0.05541 | F(2,27) = 3.2263 |
| Int, Ext, and Mult | 0.967 | 6 | 0.14642 | F(3,26) = 1.9494 |

Stim vs. Sham
| Model Noise Variables | $r^2$ | Total Free Parameters | p-value | F |
| --- | --- | --- | --- | --- |
| Base Model: | 0.949 | 3 |  |  |
| Internal Only | 0.958 | 4 | 0.86065 | F(1,28) = 0.03139 |
| External Only | 0.954 | 4 | **0.02106** | F(1,28) = 5.9753 |
| Mult Only | 0.958 | 4 | 0.10436 | F(1,28) = 2.8176 |
| Int and Ext | 0.958 | 5 | 0.07216 | F(2,27) = 2.9023 |
| Int, Ext, Mult | 0.958 | 6 | 0.18116 | F(3,26) = 1.7517 |

Post 1
| Model Noise Variables | $r^2$ | Total Free Parameters | p-value | F |
| --- | --- | --- | --- | --- |
| Base Model: | 0.956 | 3 |  |  |
| Internal Only | 0.956 | 4 | 0.95809 | F(1,28) = 0.059702 |
| External Only | 0.957 | 4 | 0.50721 | F(1,28) = 0.45134 |
| Mult Only | 0.956 | 4 | 0.66662 | F(1,28) = 0.18957 |
| Int and Ext | 0.957 | 5 | 0.80522 | F(2,27) = 0.21839 |
| Int, Ext, Mult | 0.957 | 6 | 0.92237 | F(3,26) = 0.15984 |

Post 2
| Model Noise Variables | $r^2$ | Total Free Parameters | p-value | F |
| --- | --- | --- | --- | --- |
| Base Model: | 0.979 | 3 |  |  |
| Internal Only | 0.979 | 4 | 0.58058 | F(1,28) = 0.31252 |
| External Only | 0.979 | 4 | 0.96361 | F(1,28) = 0.0021193 |
| Mult Only | 0.979 | 4 | 0.76024 | F(1,28) = 0.094962 |
| Int and Ext | 0.979 | 5 | 0.85518 | F(2,27) = 0.15735 |
| Int, Ext, Mult | 0.979 | 6 | 0.95329 | F(3,26) = 0.11033 |

**Figure 5:**
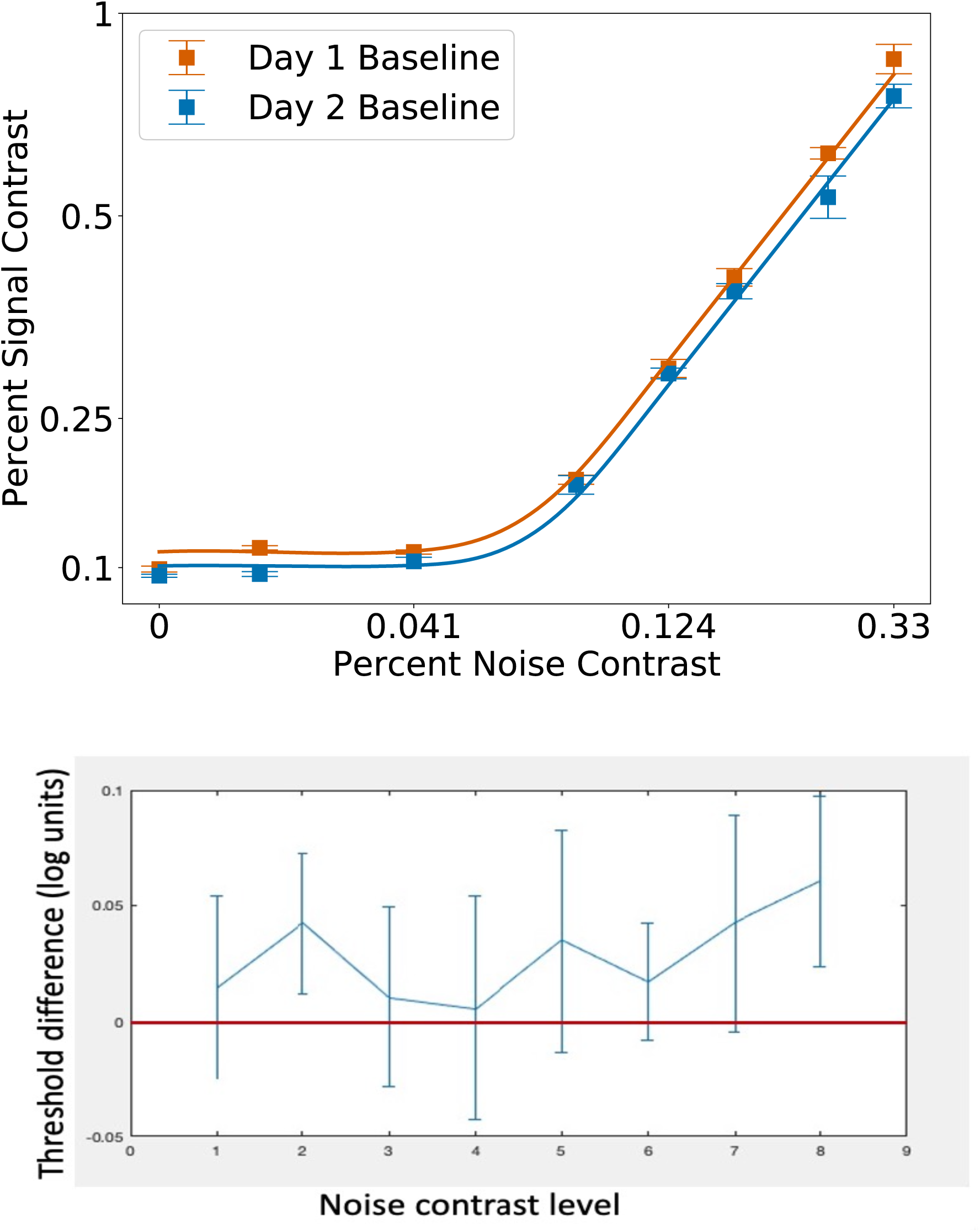
Noise signature of perceptual learning on an orientation discrimination task. (A) Fit threshold *versus* noise functions comparing Day 1 baseline (2^nd^ session) and Day 2 baseline: 79% correct thresholds are displayed. Raw data are shown as means of thresholds across subjects, with error bars as standard error of the mean. Smoothed, best-fit Perceptual Template Model curves are superimposed. (B) Difference of Day 2 baseline and Day 1 baseline (2^nd^ session) thresholds: difference of thresholds across external noise levels in log10 units. Numbers above 0 indicate improvements on Day 2 vs. Day 1.

### Online effects of tRNS

The main objective of the present study was to contrast the online effects of tRNS against those of sham stimulation. This comparison relied upon the counterbalanced, within-subject design to average out day-by-day training effects (i.e., whether sham or stimulation were received on the 2^nd^ or 3^rd^ days). More importantly, both the tRNS and the sham sessions were conducted as the second session on their respective days of testing. This controlled for any temporal effects of session order (e.g., practice and/or fatigue). Turning to the data, results shows that subjects did better at the orientation discrimination task while undergoing tRNS (Day 2/3 tRNS *versus* Day 2/3 sham), but lower contrast thresholds were only evident at high levels of external visual white noise (**Figure 6, Table 1)**—a signature of improved external noise filtering. This was confirmed with the PTM, where tRNS, relative to sham, showed a 12.0% reduction in effective external noise (*F*_*(1,28)*_*=5*.*98, p=0*.*021)*, when compared to a reduced model with fixed coefficients for additive internal noise, multiplicative internal noise, and external noise exclusion **(Figure 6)**. This improvement in external noise filtering was similar in magnitude to the above-described learning-associated changes in the same parameter (**Figure 5**). All other model parameter comparisons were not significant (**Table 1**). As with the **Figure 5** analysis, we also computed changes in performance at each noise level for visualization purposes (**Figure 6**) and found improvements to be concentrated at the higher levels of external noise.

**Figure 6:**
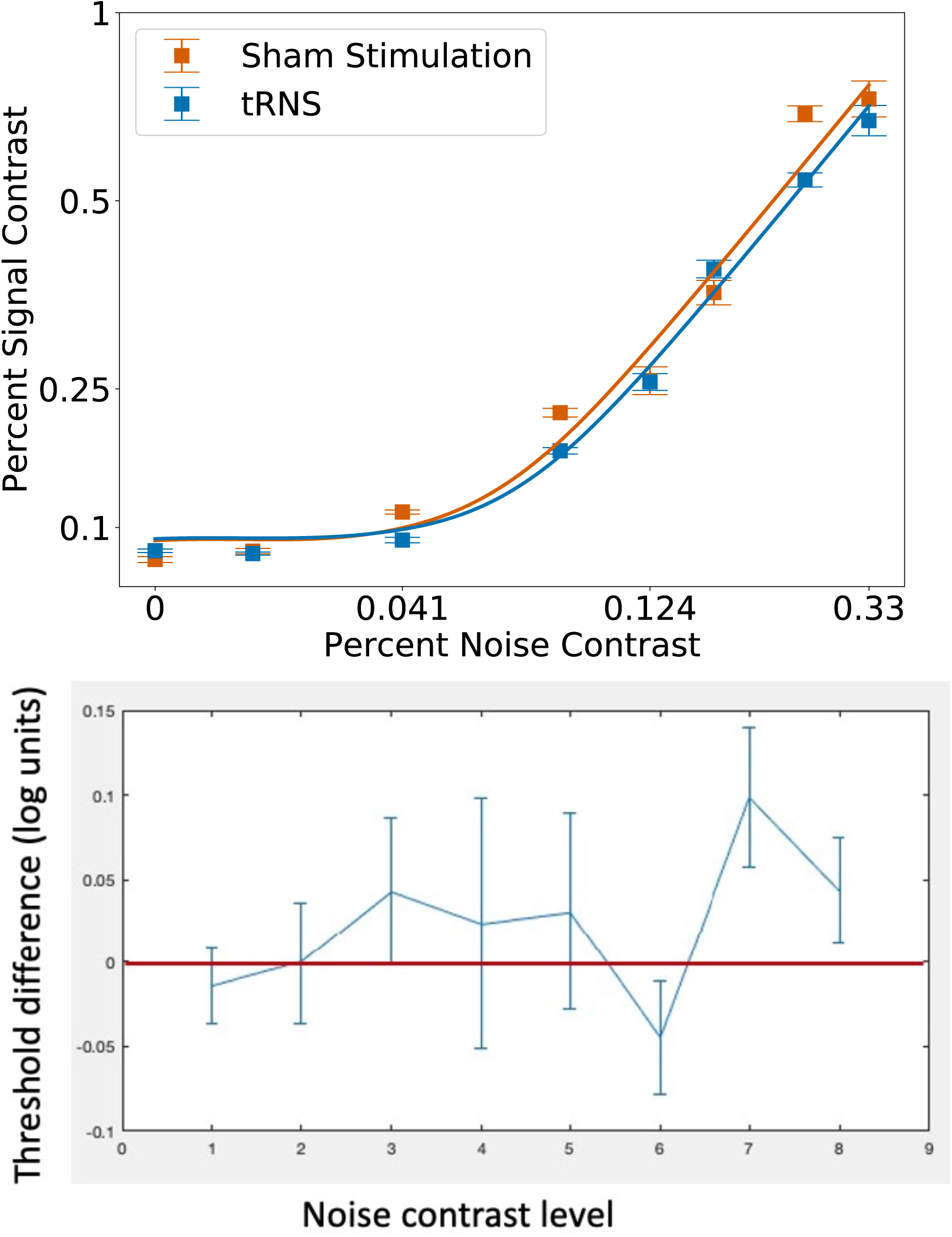
Increased External Noise Exclusion in Stimulation versus Sham condition. **(A)** Fit threshold versus noise functions comparing sham and tRNS conditions combined from days 2 and 3. 79% correct thresholds are displayed. Raw data are shown as means of thresholds across subject, error bars as standard error of the mean, with smoothed best-fit Perceptual Template Model curves superimposed. Improvements are measured only at high levels of external noise confirmed via model comparison. (B) Difference sham and thresholds: difference of thresholds across external noise levels in log10 units. Numbers above 0 indicate improvements in tRNS condition vs. Sham condition.

### Contrasting effects of tRNS within each testing session

One possible source of behavioral improvement during online stimulation is neural adaptation to tRNS-injected white noise, resulting in reduced contrast thresholds to visual white noise (see Discussion). In order to probe for this effect, we split the data from the tRNS session into two halves with the logic that if adaptation drives the observed results, these effects would be stronger in the second half of the tRNS session. We found no evidence that external noise filtering differs between the two halves (*F*_*(1,28)*_ *= 0*.*19, p = 0*.*67*). We do caution that our overall experimental design was underpowered for this split-half analysis, and it is possible that longer and/or stronger tRNS would have resulted in adaptation-like effects. However, the present findings suggest that noise adaptation was unlikely to be the major driver behind tRNS’s effects on perception.

### Offline effects of tRNS

Lastly, we assessed whether the online effects of tRNS lasted beyond the period during which stimulation was administered. To test for these offline effects, we used the same logic as in the sham *versus* tRNS analysis: we examined Post 1 sessions from Day 2 and 3, contrasting sessions that followed tRNS with those that followed sham stimulation. We performed the analogous analysis for Post 2 sessions. Thus, as we did for online tRNS analysis, these analyses compared corresponding sessions from Days 2 and 3. Here, none of the models showed significant changes (all *p>0*.*51*, **Table 1**) from a reduced model with fixed coefficients for additive internal noise, multiplicative internal noise, and external noise exclusion. Thus, we find no evidence that, with the experimental design used here, tRNS affects perceptual performance after the stimulation ends.

## Discussion

The present experiments aimed to clarify how perception of visual stimuli in noise is altered by tRNS applied bilaterally over early visual cortex in healthy, adult, human subjects. We found that tRNS caused a statistically significant boost in perception *only* when the stimuli to be discriminated contained relatively high levels of visual white noise, and *only* during application of the electrical stimulation, with no significant differences in perception after the stimulation ended.

### tRNS mimics the beneficial effects of endogenous attentional modulation

Our observation that tRNS induced a specific improvement in the ability to discriminate stimuli at high levels of external noise during task performance represents a novel insight into possible mechanism of action of this form of tES. In a classical perceptual learning experiment for which the PTM was designed (Dosher and Lu, 1998), this would be interpreted as an improvement in external noise exclusion. Improvements specifically at high levels of external noise are interpreted throughout the equivalent noise literature as improvements in the readout of sensory regions, i.e. changes occurring at a higher-level than in primary sensory areas. This is hypothesized to result from an improved higher order template that is better able to filter out the external noise from the stimulus (Dosher and Lu, 1998, Dosher and Lu, 2005). The Linear Amplifier Model (Pelli, 1981) would have referred to this change simply as increased efficiency to filter out visual white noise. Other models using the equivalent noise paradigm have also interpreted this change as more optimal (narrower) tuning to the stimulus (Ling et al., 2009, Dakin et al., 2005). All of these indicate a selective, plastic filter that allows visual cortex to better differentiate the stimulus from high amounts of visual white noise. Such improvements can result either from slow changes (hours to days) due to training-induced plasticity (as in perceptual learning), or from deployment of attentional mechanisms through selective, top-down input. We can now add tRNS to this list.

In our current result, the improvement in external noise exclusion is associated with the online application of tRNS. Compared to other results in the PTM literature, at least in its temporal properties, the effect we report here is more similar to online effects of attention rather than slow effects of perceptual learning. Moreover, in broader literature on equivalent noise tasks, an exclusive improvement at external noise exclusion is very unusual and tied to a specific type of attention. The PTM has been used to probe the effects of a variety of perceptual interventions, and amidst experiments in spatial, feature-based, transient, sustained, endogenous, and exogenous attentions (and their respective combinations) *only endogenous spatial* attention produces a shift exclusively in external noise exclusion (Dosher and Lu, 1999, Lu and Dosher, 1999, Dosher and Lu, 2000, Dosher and Lu, 2005). Endogenous spatial attention is characterized by slightly slower deployment (when compared to exogenous attention), voluntary control, and its application to specific areas of the visual field (Carrasco, 2011). This type of attention is well established to improve perceptual performance on a variety of tasks, including the specific orientation discrimination performed in this study (Dosher and Lu, 2000) and is primarily associated with activity in a dorsal frontoparietal network (Carrasco, 2011, Corbetta and Shulman, 2002). It is also worth noting that both tRNS and endogenous spatial attention are known to enhance visual perceptual learning (Herpich et al., 2019, Donovan and Carrasco, 2018).

While the effect of tRNS strongly resembles the *effect* of endogenous attention, we do not argue that tRNS *directly* stimulates or induces endogenous attention. Firstly, while studies using tRNS to boost attentional effects have produced positive results akin to those found in visual perception (Tyler et al., 2018), these studies were performed with electrode configurations over parietal and frontal cortices – i.e., neural loci well-established for the origination and processing of attentional signals. Given that our stimulation sites were directly over occipital cortex, it is extremely unlikely that the observed perceptual enhancements were caused by directly stimulated endogenous spatial attention. Secondly, a bottom-up excitatory attention-like signal resulting from tRNS would function more like added salience to the stimulus or stimulus location. However, if salience were the improvement mechanism for tRNS in perceptual processing, our results should resemble those of *exogenous* attentional mechanisms on threshold *versus* noise curves. Instead of a boost in salience, our effect instead resembles the internal generation of spatial attention, in spite of – as noted above – lack of direct stimulation of neural areas associated with the generation of attention.

### tRNS and Visual Noise

Recent work showed that the effects of tRNS can mimic those of visual white noise (van der Groen et al., 2019, van der Groen et al., 2018, van der Groen and Wenderoth, 2016). The central role of noise in the paradigm used here raises a number of questions. Namely, if the visual system treats tRNS as visual white noise, why do thresholds improve rather than worsen in the presence of tRNS, and why do thresholds improve only at high levels of external noise? In the PTM and related models, the internal noise is assumed to include a wide array of signal transmission, neural coding, and perceptual inefficiencies inherent in the visual system (Lu and Dosher, 2008). Furthermore, the PTM hinges on the logic that by exceeding the quantity of internal noise using additional, external, visual noise, we can estimate the quantity of this amalgam of internal noise. Our results present two hypotheses about tRNS in relation to internal noise in this orientation discrimination task. Firs, we can assume that if the visual system treats tRNS as visual white noise (van der Groen et al., 2019, van der Groen et al., 2018, van der Groen and Wenderoth, 2016), the *quantity* of this tRNS white noise does not exceed the quantity of noise estimated as internal noise. If it did, we would expect to see a change in performance at low levels of external noise, which we do not. Alternatively, the noise added to the visual system due to tRNS may not be measured by the internal noise in the PTM at all. If it were, we would expect a baseline shift in the quantity of internal noise when tRNS was switched on. Given that we found no effects of tRNS at low levels of internal noise, the parsimonious conclusion is that the internal noise measurement does not include additional noise from tRNS. However, the possibility remains that another, unknown, beneficial effect of tRNS balances out the negative effects on “brain” noise caused by tRNS.

It is also unlikely that our observed effects of tRNS can be explained using the proposed theory of stochastic resonance (Fertonani and Miniussi, 2017, van der Groen and Wenderoth, 2016), a mechanism where noninformative white noise boosts subthreshold stimuli above threshold (Gammaitoni et al., 1989). van der Groen and Wenderoth (2016) elegantly showed that systematically increasing the current amplitude of tRNS during a visual discrimination produces an inverted U-shaped curve typical of stochastic resonance with the best performance (lower thresholds) at 1mA tRNS amplitude. The authors find similar results by measuring performance on the same task under increasing amounts of <u>visual</u> white noise, again demonstrating stochastic resonance. However, it is not clear that stochastic resonance applies to typical tRNS stimulation parameters, such as those used in our study. Our stimulation amplitude of 2mA is higher than the highest amplitude tested by van der Groen and Wenderoth (2016), who found that effects of stochastic resonance disappeared by 1.5mA. Moreover, van der Groen & Wenderoth utilized stimulus-locked short bursts of online tRNS (2.04s at a time), whereas most tRNS studies utilize longer blocks of stimulation (> 10 min), similar to what was employed here - e.g. (Herpich et al., 2019, Fertonani et al., 2011, Pirulli et al., 2013). This is relevant, as it was demonstrated by Chaieb et al (2011) that tRNS stimulation lasting 5 minutes (or longer) increases cortical excitability in motor cortex (assessed by TMS-induced MEPs), suggesting that van der Groen & Wenderoth’s effect may rely on a short-term, induced white noise signal rather than a change in excitability. In sum, it is possible and perhaps likely that the mechanisms of tRNS action vary significantly with stimulation parameters. This motivated our work targeting tRNS mechanisms but using stimulation parameters that are more typically used in brain modulation studies. Nontheless, it is worth noting that our results do not rule out the possibility that tRNS is injecting the equivalent of white noise into the visual system during stimulation. However, this account of tRNS’s effects does lead to specific predictions. Namely, if tRNS acts on the brain as white noise (van der Groen and Wenderoth, 2016), then there should be a point at which tRNS *impairs* performance. This point, and its defining parameter space, remain to be determined.

Finally, we considered the possibility that tRNS improves performance by reducing the effects of visual noise used in our paradigm. Specifically, the brain might have adapted to tRNS-induced white noise, reducing the effective contrast of external white noise used in our study. Neural adaptation is a well-studied, canonical principle of visual processing (Carandini and Heeger, 2012). The main consequence of adaptation is reduced sensitivity to the adapted feature, a result that is found for a wide range of visual stimuli and timescales of adaptation (Barlow and Hill, 1963, Kohn, 2007, Mather et al., 1998, Webster, 2011, Glasser et al., 2011). Thus, if tRNS acts as a form of white noise that is sufficiently similar to visual noise, then sustained delivery of tRNS could result in adaptation that reduces the impact of the visual noise on vision (e.g., 50% contrast visual noise now behaves like 40% contrast noise). If this were true, we would expect a shift in the threshold *versus* contrast curve to the right. The effects of such a shift would be most prominent at high levels of noise, matching the external noise reduction results we report here. To test this hypothesis, we performed a split-half analysis on the 1^st^ and 2^nd^ halves of the tRNS-stimulation sessions. While, as noted in the Results section, our analysis may be underpowered for a clear indication of this effect, we saw no evidence indicating that adaptation to noise was a likely basis for our main result.

### Practical utility of equivalent noise in studying transcranial electrical stimulation

One of the critical and challenging issues present in the literature surrounding all transcranial stimulation is the sheer size of parameter space that an experimenter must navigate in order to successfully achieve an effect. That is, choosing stimulation current (generally between 0 and 3mA for safety (Bikson et al., 2016), type (tDCS, tRNS, tACS), electrode configuration (unilateral stimulation, bilateral stimulation, stimulation of higher order decision making areas *versus* primary sensory regions, etc.), stimulation timing (blocks of 10 minute or 20 minute stimulation, epochs of 4 minutes on and 2 minutes off (Fertonani et al., 2011), etc., and stimulation and task timing (online vs offline). One of the common outcomes of this impractically large parameter space is that conflicting results abound (reviewed in Nitsche et al. (2008), Fertonani and Miniussi (2017)). The equivalent noise approach used presently allowed us to explain why this might be the case and enabled us to illustrate the fact that, had we performed this experiment using an identical task and identical stimuli, but without noise, or only low levels of noise, we would have reported a null result for the effect of tRNS.

## Conclusions

tRNS-induced behavioral improvements in visual orientation discrimination were only observed during online stimulation and for visual targets containing high levels of external visual noise. While the exact mechanisms of this effect remain unclear, the evidence presented here supports recent findings that tRNS acts on sensory cortex in a manner that modestly, but specifically, improves perceptual performance <u>during</u> stimulation. These effects mimic known effects of endogenous spatial attention. Furthermore, our data help to understand many instances of null results in the use of tRNS by showing how they can arise when stimulus noise is low or absent. We hope that future work will continue to elucidate mechanisms underlying behavioral effects of tRNS, with similarities between endogenous spatial attention and tRNS reported here offering a worthwhile direction to pursue.

## Author Contributions

MDM designed, programmed, piloted, conducted, analyzed, and wrote the manuscript. WJP programmed stimulus display and analysis code, advised on analysis and edited the manuscript. SK piloted and ran subjects. SC piloted and ran subjects. LB provided stimulation parameters and training, and edited the manuscript. AB piloted and tested stimulation parameters, and edited the manuscript. KRH designed, analyzed and edited the manuscript. DT designed, analyzed and edited the manuscript.

## Acknowledgments

We thank Drs. Ruyuan Zhang and Oh-Sang Kwon for experimental design and analysis methodologies relating to presentation and fitting of the Perceptual Template Model. We thank the Tadin and Huxlin lab members for ongoing feedback on experimental results and interpretation of data.

## Data Availability Statement

All experimental data is available from the corresponding authors at request via email in the form of labeled MATLAB exports.

